# A Mental Health Paradox: Mental health was both a motivator and barrier to physical activity during the COVID-19 pandemic

**DOI:** 10.1101/2020.09.03.280719

**Authors:** Maryam Marashi, Emma Nicholson, Michelle Ogrodnik, Barbara Fenesi, Jennifer Heisz

## Abstract

The COVID-19 pandemic has impacted the mental health, physical activity, and sedentary behavior of citizens worldwide. Using an online survey with 1669 respondents, we sought to understand why and how by querying about perceived barriers and motivators to physical activity that changed because of the pandemic, and how those changes impacted mental health. Consistent with prior reports, our respondents were less physically active (aerobic activity, −11%, *p* <0.05; strength-based activity, −30%, *p*<0.01) and more sedentary (+11%, *p*<0.01) during the pandemic as compared to 6-months before. The pandemic also increased psychological stress (+22%, *p* <0.01) and brought on moderate symptoms of anxiety and depression. Respondents’ whose mental health deteriorated the most were also the ones who were least active (depression *r* = −.21, *p*<0.01; anxiety *r* = −.12, *p*<0.01). The majority of respondents were unmotivated to exercise because they were too anxious (+8%, *p* <0.01), lacked social support (+6%, *p* =<0.01), or had limited access to equipment (+23%, *p* <0.01) or space (+41%, *p* <0.01). The respondents who were able to stay active reported feeling less motivated by physical health outcomes such as weight loss (−7%, *p*<0.01) or strength (−14%, *p*<0.01) and instead more motivated by mental health outcomes such as anxiety relief (+14%, *p* <0.01). Coupled with previous work demonstrating a direct relationship between mental health and physical activity, these results highlight the potential protective effect of physical activity on mental health and point to the need for psychological support to overcome perceived barriers so that people can continue to be physically active during stressful times like the pandemic.

## Introduction

During the initial phase of the COVID-19 pandemic, governing bodies worldwide took decisive action to protect their citizens against the novel coronavirus by enforcing public lockdown and closing all non-essential services [1]. Although such measures helped to “flatten the curve” and minimize infection rates, the restrictions also had unforeseen consequences on citizens’ health and well-being in that pandemic-related concerns amplified mental distress of citizens worldwide [2–8]. A major concern is that psychological distress can quickly deteriorate into mental illness, even for people without a prior diagnosis [9]; though susceptibility varies by age [10] and income [11], with younger and low income being more susceptible. This has created an urgent need for effective interventions to help minimize the psychological burden of the pandemic and prevent a mental health crisis [12].

One of the most effective interventions to prevent stress-induced mental illness is physical activity. People who are more active also tend to be less anxious and depressed [13], and sedentary people who engage in a new exercise program experience relief from their depressive symptoms [14]. Compared to antidepressant medication, thirty minutes of *moderate-intensity* aerobic exercise three times weekly may be equally effective at reducing psychological distress and decreasing symptoms of depression and anxiety without any of the drug-related side effects such as nausea, fatigue, or loss of appetite [15].

However, maintaining a regular exercise program is difficult at the best of times and the conditions surrounding the COVID-19 pandemic may be making it even more difficult. The World Health Organization (WHO) recommends adults participate in 150 min/week of moderate-intensity aerobic physical activity (or 75 min/week of vigorous-intensity aerobic physical activity) and 2 or more days per week of muscle-strengthening activities [16]. Globally, about 1 in 4 people were not meeting these guidelines prior to the pandemic [16], with these numbers differing by age [17] and income [18] such that younger adults are more likely to meet guidelines than older adults and higher income predicts better adherence to the guidelines than lower income. Recent reports suggest the pandemic has further decreased physical activity and increased sedentary time [19]. In animal models, forced inactivity causes depressive symptoms [20], and experimentally controlled periods of exercise withdrawal in humans lead to increased symptoms of depression and anxiety [21]. This link between physical and mental health is being further exposed in humans during this pandemic. An online survey administered during the initial stage of the COVID-19 pandemic found that respondents who were less physically active had worse mental health [22]. Another survey conducted at the same time found that more screen time (a common sedentary behaviour) was associated with worse mental health in all respondents except for those who were physically active [23], suggesting that physical activity may protect against the expected mental health decline caused by the sedentary lifestyle of enforced lockdown.

What is it about the pandemic that is making people less active, and how can we best support those who are struggling to stay active? Using an online survey, the present study gathered information from 1669 respondents pertaining to their physical activity, sedentary behavior and mental health before and during the initial lockdown of COVID-19. We were primarily interested in investigating any shifts in perceived barriers and motivators to being physically active. Typically, the most common perceived barrier to being physically active is lack of time [24]. Lack of motivation is another commonly cited barrier [24]. However, given the unprecedented circumstances surrounding the COVID-19 pandemic, people may now face unique barriers and motivators to engaging in physical activity.

## Materials and methods

### Design and respondents

To achieve a small margin of error based on a population size of 37 million, we recruited a total of 1669 respondents over a two-month data collection period (April 23 to June 30, 2020), for a 2% margin of error with 95% confidence intervals. The survey was open to all respondents at least 18 years of age, fluent in English, and able to complete the online survey. Respondents were recruited through the personal social media accounts of the research team and through local news sources (news articles by media at McMaster University and Hamilton Spectator). Respondents were also recruited via a link provided at the end of an op-ed piece published in The Conversation Canada, a national independent news source from the academic and research community.

The survey consisted of 30 questions and used a mix of multiple-choice, single choice, and short answer questions to query respondents about their demographic information, and their current and past (prior to the pandemic) physical activity behaviour (minutes/week). Additionally, respondents were asked about their current and past mental health status (i.e., stress levels, anxiety and depressive symptoms). All questions pertaining to physical activity and mental health were designed using validated rating scales. Respondents were included in a draw for 20 cash prizes of $100 CAN as remuneration for their participation in the form of an emailed prepaid voucher.

### Measurements

#### Physical activity

The Physical Activity and Sedentary Behavior Questionnaire (PASB-Q) [25] was adapted (i.e., rewording of questions to include COVID-19) to quantify self-reported levels of physical activity and sedentary behaviour 6-months prior to and during the COVID-19 pandemic. Respondents were asked to report minutes/week of strength training and aerobic exercise, hours/week of sedentary behavior, and self-rated activity level status on a 5-point scale where 1 = “Completely sedentary”, 2 = “Slightly active”, 3 = “Very active”, 4 = “Recreational athlete”, 5 = “Elite athlete”.

#### Barriers and motivators to exercise

Respondents were asked to report current and prior (i.e., 6 months prior to COVID-19) barriers preventing them from being physically active using a multiple-choice list (e.g., “I could/cannot find the time in my day”, “I did/do not have access to a gym or recreational facility”) and motivators encouraging them to be physically active (e.g., “To maintain a healthy body weight”, “To build muscle and/or strength”) (S1 Appendix).

#### Mental health

Anxiety was measured using an adapted (i.e., on a 5-point scale instead of a 3-point scale to match other questionnaires and ease participant burden) version of the Generalized Anxiety Disorder 7-item Scale (GAD-7) [26]. Respondents were asked how often they felt bothered by each anxiety symptom since the onset of COVID-19. Response options were 1 = “Not at all”, 2 = “Several days”, 3 = “More than half the days”, 4 = “Most days”, and 5= “Every day”. All seven items were combined to form a global measure of anxiety.

Depression was measured using an adapted version of the Patient Health Questionnaire (PHQ-9) [27]; all but one of the 9 items (i.e., the one pertaining to suicidal thoughts and/or self-harm) were included for a total of 8 items, which were combined into a global measure of depression. Respondents were asked how often they feel bothered by each depression symptom since the onset of COVID-19. Response options were 1= “Not at all”, 2 = “Several days”, 3 = “More than half the days”, 4 = “Most days”, and 5 = “Every day”.

Question 3 from the Perceived Stress Scale (PSS) [28] was used to measure psychological stress. Respondents were asked how often they felt nervous and “stressed” both prior to and since the onset of COVID-19 on a 5-point scale where 1 = “Never”, 2 = “Sometimes”, 3 = “Fairly often”, 4 = “Often”, and 5 = “Very often”.

To capture an overall change in mental health since the onset of COVID-19, respondents were asked to rate their overall mental health since COVID in relation to how it was in the six-months prior to COVID with the options of choosing “Much better”, “Better”, “No change”, “Worse”, or “Much worse”.

#### Statistical Analyses

The IBM SPSS® statistics software platform (Version 26) was used to carry out all analyses. Descriptive statistics (means and standard deviations for continuous variables, and frequency counts and percentages for categorical variables) were computed to describe demographic characteristics, mental health, and physical activity levels. Normality was assessed using Shapiro-Wilkes tests and through visual inspection of histograms. For all analyses, significance was considered at *p* < 0.05, and nonparametric tests were chosen wherever data did not meet the assumption of normality.

For correlational analysis, all respondents who left 100% of the survey questions blank were removed (N=166). Physical activity and mental health data were then screened for missingness which ranged from 8.2-11.8% and 10.2-17.3% respectively. Missing cells were subsequently imputed using expectation-maximization [29] for all physical activity and mental health variables. In the case where a negative physical activity datum or score exceeding the maximum mental health score was imputed, the datum was removed. Physical activity and mental health data used in correlations had a resulting 0.1-0.5% and 0.1% missingness respectively.

## Results

### Sample characteristics and mental health status

Survey respondents were primarily female between 18-29 years of age, living in Canada and well-educated (Table 1). Most respondents spent at least four weeks in social isolation at the time of the survey, and a large portion was currently working regular hours from home. More respondents reported that they were making “less than enough” since the onset of the pandemic compared to their income within the 6 months before the pandemic. Although few respondents indicated a close exposure to someone with COVID-19 or COVID-19 symptoms, nearly half knew someone immunocompromised and therefore at high risk.

**Table 1.**
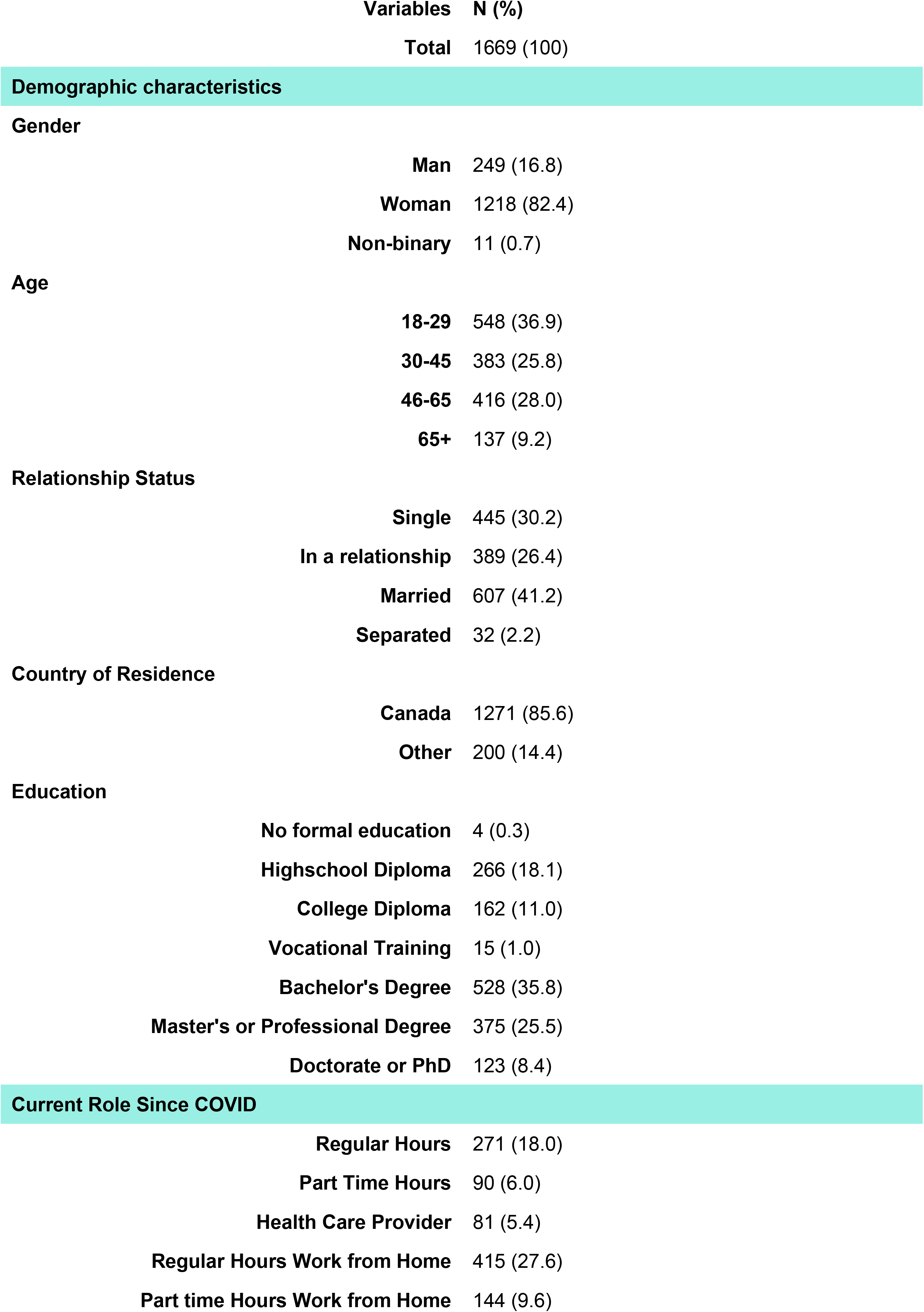
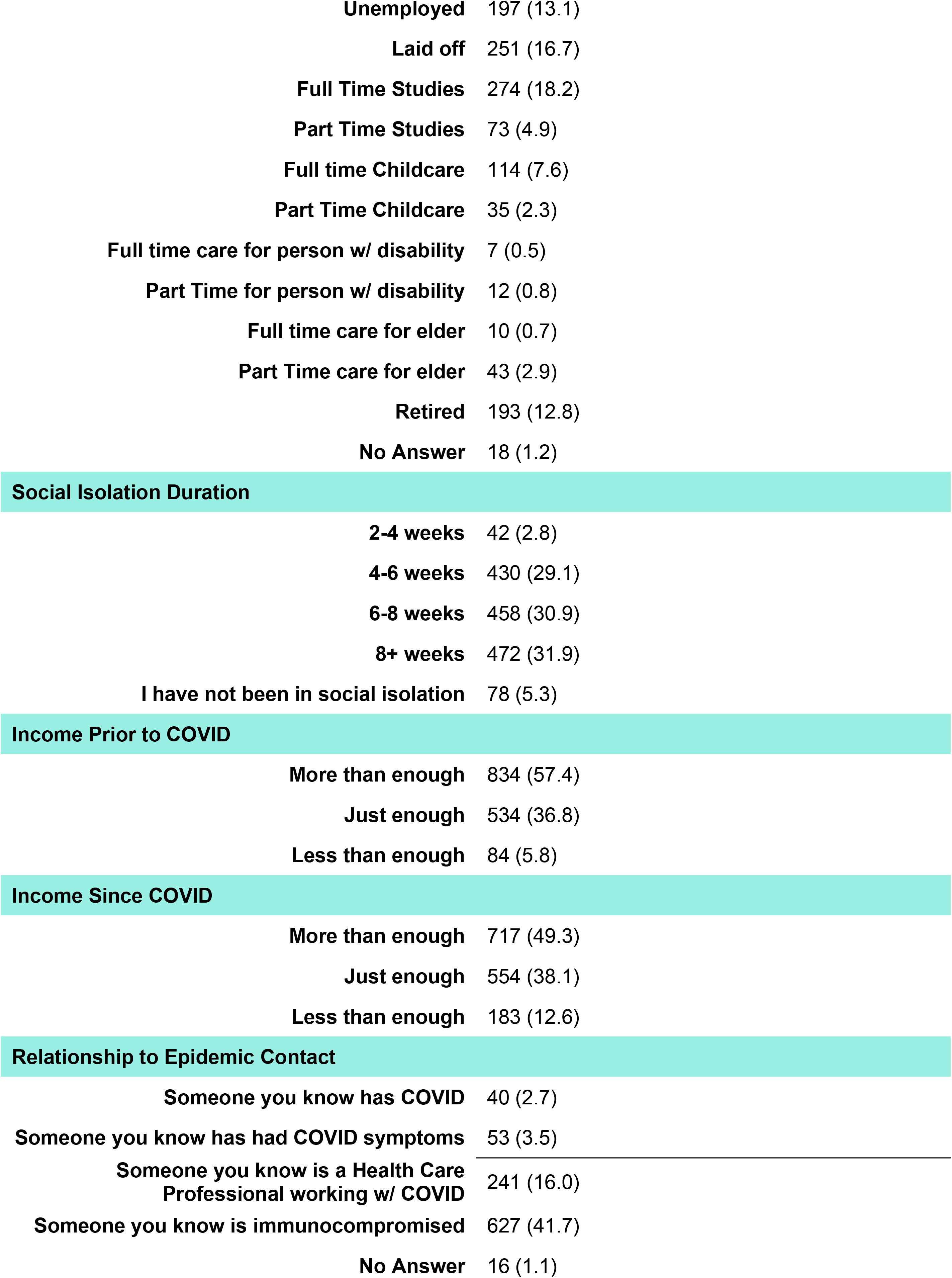
Sample Description

| Variables |  | N (%) |
| --- | --- | --- |
| Total |  | 1669 (100) |
| Demographic characteristics |  |  |
| Gender |  |  |
|  | Man | 249 (16.8) |
|  | Woman | 1218 (82.4) |
|  | Non-binary | 11 (0.7) |
| Age |  |  |
|  | 18-29 | 548 (36.9) |
|  | 30-45 | 383 (25.8) |
|  | 46-65 | 416 (28.0) |
|  | 65+ | 137 (9.2) |
| Relationship Status |  |  |
|  | Single | 445 (30.2) |
|  | In a relationship | 389 (26.4) |
|  | Married | 607 (41.2) |
|  | Separated | 32 (2.2) |
| Country of Residence |  |  |
|  | Canada | 1271 (85.6) |
|  | Other | 200 (14.4) |
| Education |  |  |
|  | No formal education | 4 (0.3) |
|  | Highschool Diploma | 266 (18.1) |
|  | College Diploma | 162 (11.0) |
|  | Vocational Training | 15 (1.0) |
|  | Bachelor's Degree | 528 (35.8) |
|  | Master's or Professional Degree | 375 (25.5) |
|  | Doctorate or PhD | 123 (8.4) |
| Current Role Since COVID |  |  |
|  | Regular Hours | 271 (18.0) |
|  | Part Time Hours | 90 (6.0) |
|  | Health Care Provider | 81 (5.4) |
|  | Regular Hours Work from Home | 415 (27.6) |
|  | Part time Hours Work from Home | 144 (9.6) |

|  |  |
| --- | --- |
| **Unemployed** | 197 (13.1) |
| **Laid off** | 251 (16.7) |
| **Full Time Studies** | 274 (18.2) |
| **Part Time Studies** | 73 (4.9) |
| **Full time Childcare** | 114 (7.6) |
| **Part Time Childcare** | 35 (2.3) |
| **Full time care for person w/ disability** | 7 (0.5) |
| **Part Time for person w/ disability** | 12 (0.8) |
| **Full time care for elder** | 10 (0.7) |
| **Part Time care for elder** | 43 (2.9) |
| **Retired** | 193 (12.8) |
| **No Answer** | 18 (1.2) |

Social Isolation Duration
|  |  |
| --- | --- |
| **2-4 weeks** | 42 (2.8) |
| **4-6 weeks** | 430 (29.1) |
| **6-8 weeks** | 458 (30.9) |
| **8+ weeks** | 472 (31.9) |
| **I have not been in social isolation** | 78 (5.3) |

Income Prior to COVID
|  |  |
| --- | --- |
| **More than enough** | 834 (57.4) |
| **Just enough** | 534 (36.8) |
| **Less than enough** | 84 (5.8) |

Income Since COVID
|  |  |
| --- | --- |
| **More than enough** | 717 (49.3) |
| **Just enough** | 554 (38.1) |
| **Less than enough** | 183 (12.6) |

Relationship to Epidemic Contact
|  |  |
| --- | --- |
| **Someone you know has COVID** | 40 (2.7) |
| **Someone you know has had COVID symptoms** | 53 (3.5) |
| **Someone you know is a Health Care Professional working w/ COVID** | 241 (16.0) |
| **Someone you know is immunocompromised** | 627 (41.7) |
| **No Answer** | 16 (1.1) |

### Impact of the pandemic on mental health status

Average anxiety (19.0±0.2, max = 35) and depression (20.2±0.2, max = 40) scores reflect moderate symptoms for both anxiety and depression. Tables 2 and 3 show positive correlations between anxiety and depression from the onset of the pandemic revealing that individuals with more anxiety symptoms also had more depressive symptoms. To identify respondents at higher risk for mental illness during the pandemic, Kruskal-Wallis one-way ANOVA evaluated between-group differences in anxiety and depression by age and income. With respect to age, respondents aged 18-29 experienced significantly higher anxiety than those 30-45 (*H*(4,1310)= 2.85, *p* = < 0.01), 46-65 (*H*(4,1310)= 4.15, *p* = < 0.01), and 65+ (*H*(4,1310)= 7.85, *p* = < 0.01). The same pattern was seen in depression, wherein those aged 18-29 experienced higher depression than those 30-45 (*H*(4,1278)= 5.13, *p* = < 0.01), 46-65 (*H*(4,1278)= 5.92, *p* = < 0.01), and 65+ (*H*(4,1278)= 7.61, *p* = < 0.01). With respect to income, respondents making “less than enough” had significantly higher levels of anxiety and depression than those making “just enough” (H(3,1310) = 6.93, *p* < 0.01; H(3,1278) = 9.19, *p* < 0.01) and “more than enough” (H(3,1310) = 8.64, *p* < 0.01; H(3,1278) = 10.96, *p* < 0.01). As well, respondents making “just enough” had significantly higher anxiety and depression than those making “more than enough” (H(3,1310) = −3.71 *p* < 0.01; H(3,1278) = −4.40, *p* < 0.01).

**Table 2.**
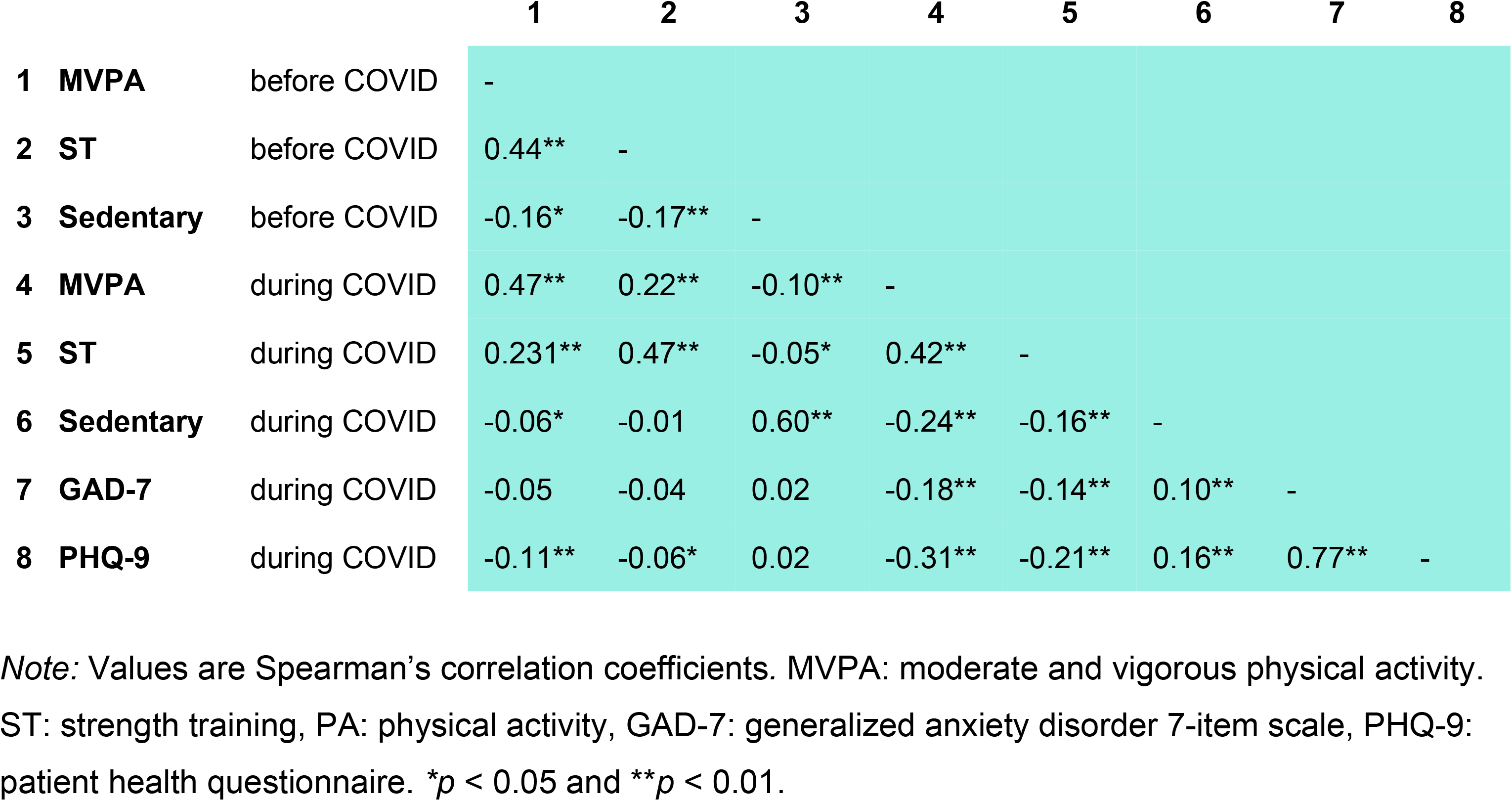
Correlations between physical activity, sedentary behaviour and mental health

|  |  | 1 | 2 | 3 | 4 | 5 | 6 | 7 | 8 |
| --- | --- | --- | --- | --- | --- | --- | --- | --- | --- |
| 1 | MVPA before COVID | - |  |  |  |  |  |  |  |
| 2 | ST before COVID | 0.44** | - |  |  |  |  |  |  |
| 3 | Sedentary before COVID | -0.16* | -0.17** | - |  |  |  |  |  |
| 4 | MVPA during COVID | 0.47** | 0.22** | -0.10** | - |  |  |  |  |
| 5 | ST during COVID | 0.231** | 0.47** | -0.05* | 0.42** | - |  |  |  |
| 6 | Sedentary during COVID | -0.06* | -0.01 | 0.60** | -0.24** | -0.16** | - |  |  |
| 7 | GAD-7 during COVID | -0.05 | -0.04 | 0.02 | -0.18** | -0.14** | 0.10** | - |  |
| 8 | PHQ-9 during COVID | -0.11** | -0.06* | 0.02 | -0.31** | -0.21** | 0.16** | 0.77** | - |

*Note:* Values are Spearman's correlation coefficients. MVPA: moderate and vigorous physical activity. ST: strength training, PA: physical activity, GAD-7: generalized anxiety disorder 7-item scale, PHQ-9: patient health questionnaire. \* $p < 0.05$ and \*\* $p < 0.01$ .

**Table 3.**
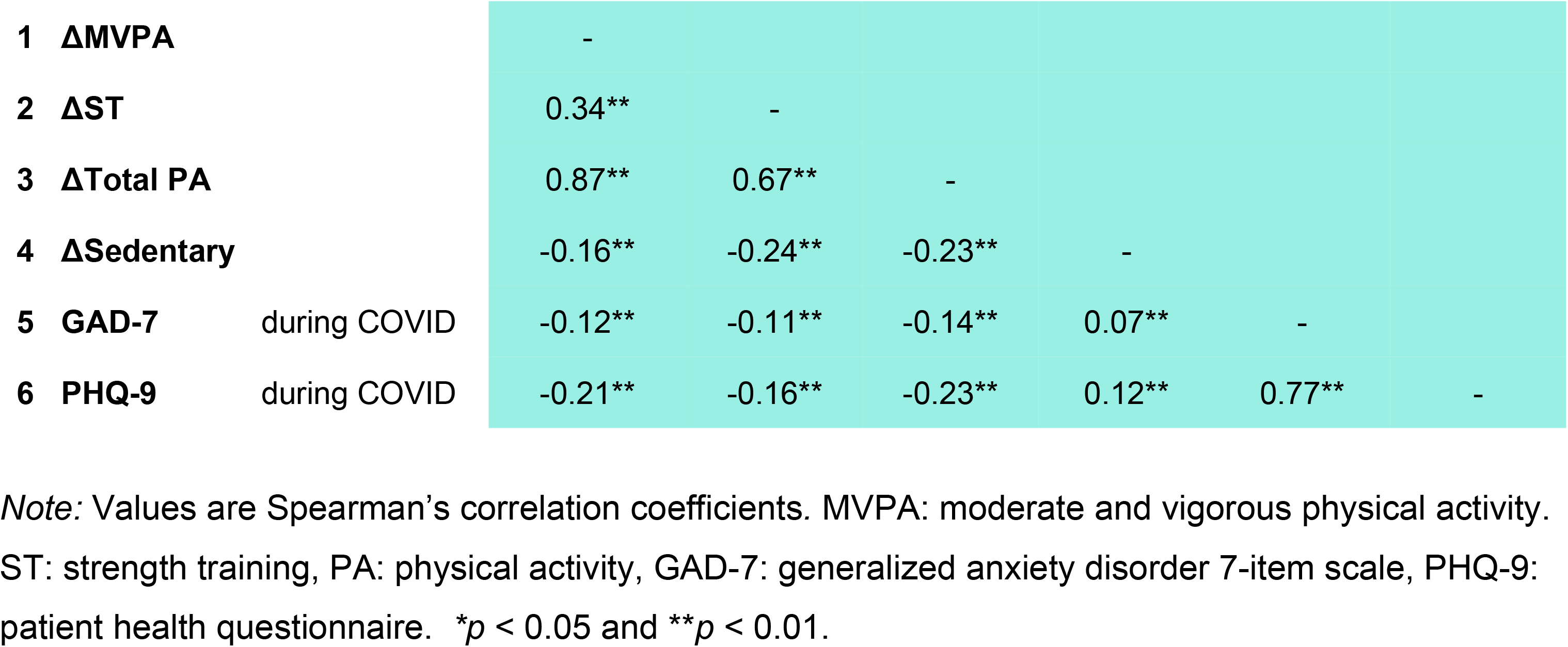
Correlations between changes in physical activity and current mental health status.

To assess changes in self-perceived psychological stress, a Wilcoxon Signed Ranks Test was performed on ratings before and during the initial stages of the COVID-19 pandemic and McNemar’s tests were used to assess changes in frequencies. There was a significant increase in stress levels during the pandemic (*Z* = −17.00, *p* < 0.01) (Figure 1). Since the onset of COVID-19, 22% of respondents who had felt stressed “sometimes” (*p* < 0.01) now felt stressed “often” (+7%; *p* < 0.01) or “very often” (+17%; *p* < 0.01) (Figure 2). The pandemic did not impact the number of respondents who reported “never” (*p* = 0.13) feeling stressed or feeling stressed “fairly often” (*p* = 0.28).

**Fig 1.**
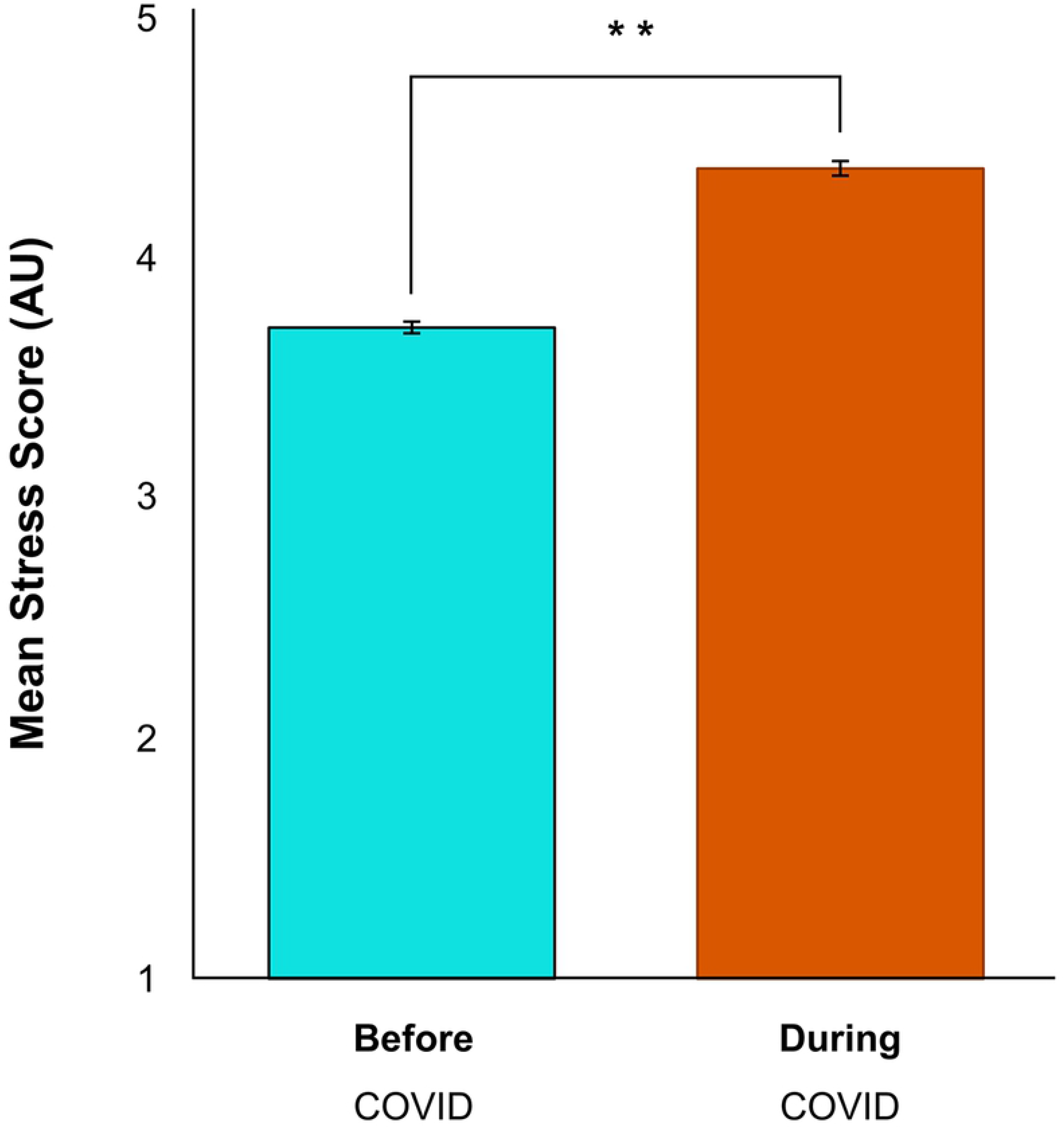
Psychological stress increased significantly during the COVID-19 pandemic (\*\**p* < 0.01). Error bars represent standard error.

**Fig 2.**
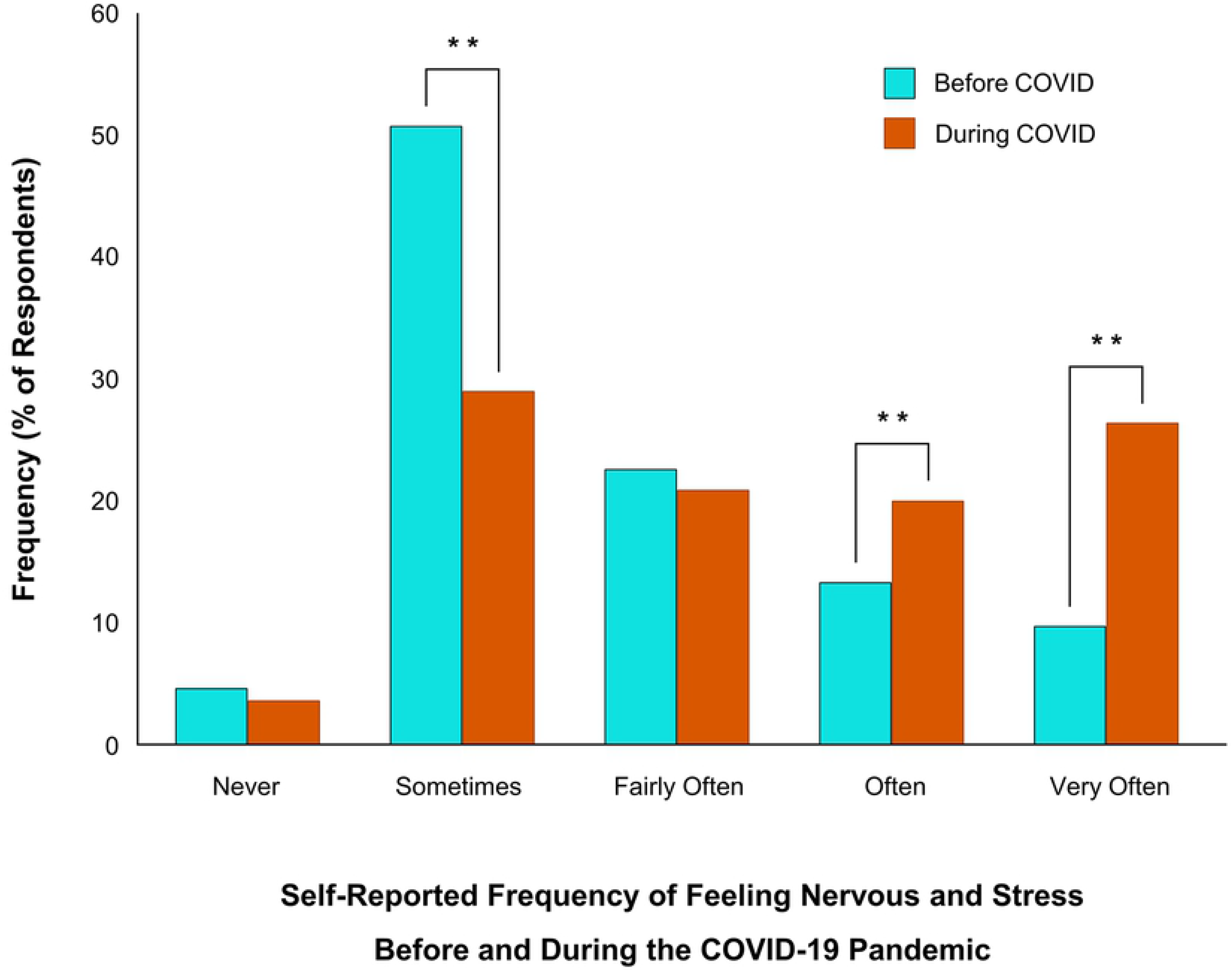
Changes in self-report psychological stress before and during the COVID-19 pandemic. 55% of respondents indicated their overall mental health had gotten “worse” or “much worse” during the COVID-19 pandemic (\*\**p* < 0.01).

### Impact of the pandemic on physical activity and sedentary behaviour

To test the hypothesis that physical activity levels dropped during the initial stages of the COVID-19 pandemic, Wilcoxon Signed Rank statistics were computed on changes in aerobic activity, strength training activity and sedentary behaviour and McNemar’s tests were used to assess the change in self-identified exercise status. Since the onset of COVID-19, respondents’ aerobic activity decreased by 22 minutes (−11%; Z= −2.50, *p* < 0.05), their strength-based activity decreased by 32 minutes (−30%; Z= −7.89, *p* < 0.01), and their sedentary times increased by 33 minutes (+11%; *Z* = −14.18, *p* < 0.01) (Figures 3 and 4). Respondents who had been “recreational athletes” (−6%; *p* < 0.01), “very active” (−6%; *p* < 0.01), or “moderately active” (−5%; *p* < 0.01) before the pandemic, now identify as being “completely sedentary” (+17%; *p* < 0.01). There was no change in the frequency of respondents who self-identified as “elite athletes” (*p* = 0.12) (Figure 5). Tables 2 and 3 show correlations between physical activity and sedentary behaviour both before and during the COVID-19 pandemic. Although total physical activity decreased during COVID-19, each respondents’ physical activity level remained proportional to their activity level prior to the pandemic.

**Fig 3.**
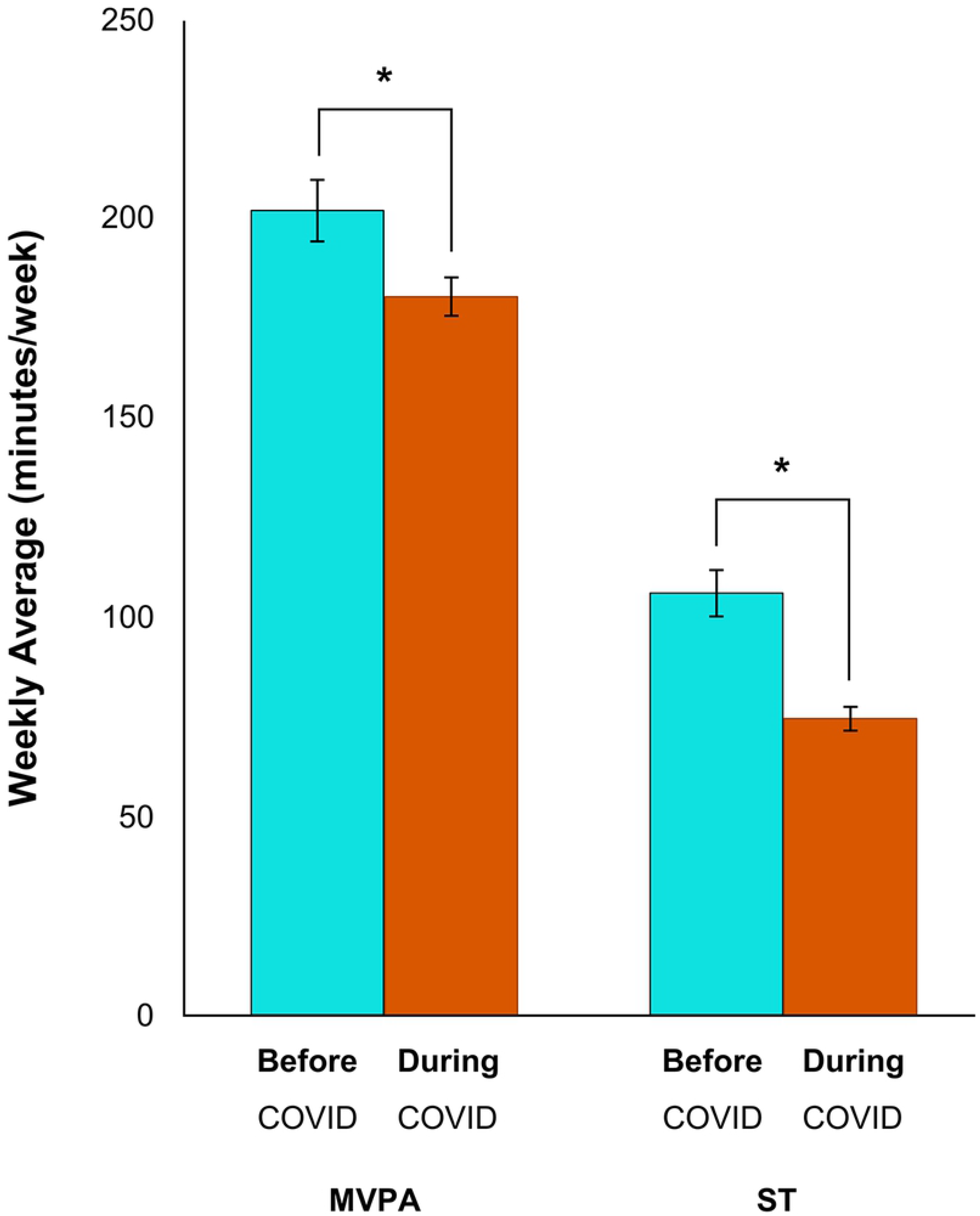
Changes in physical activity levels before and during the COVID-19 pandemic. There was a significant decrease in both moderate-to-vigorous physical aerobic activity (MVPA; \**p* < 0.05) and strength training (ST; \*\**p* < 0.01). Error bars represent standard error.

**Fig 4.**
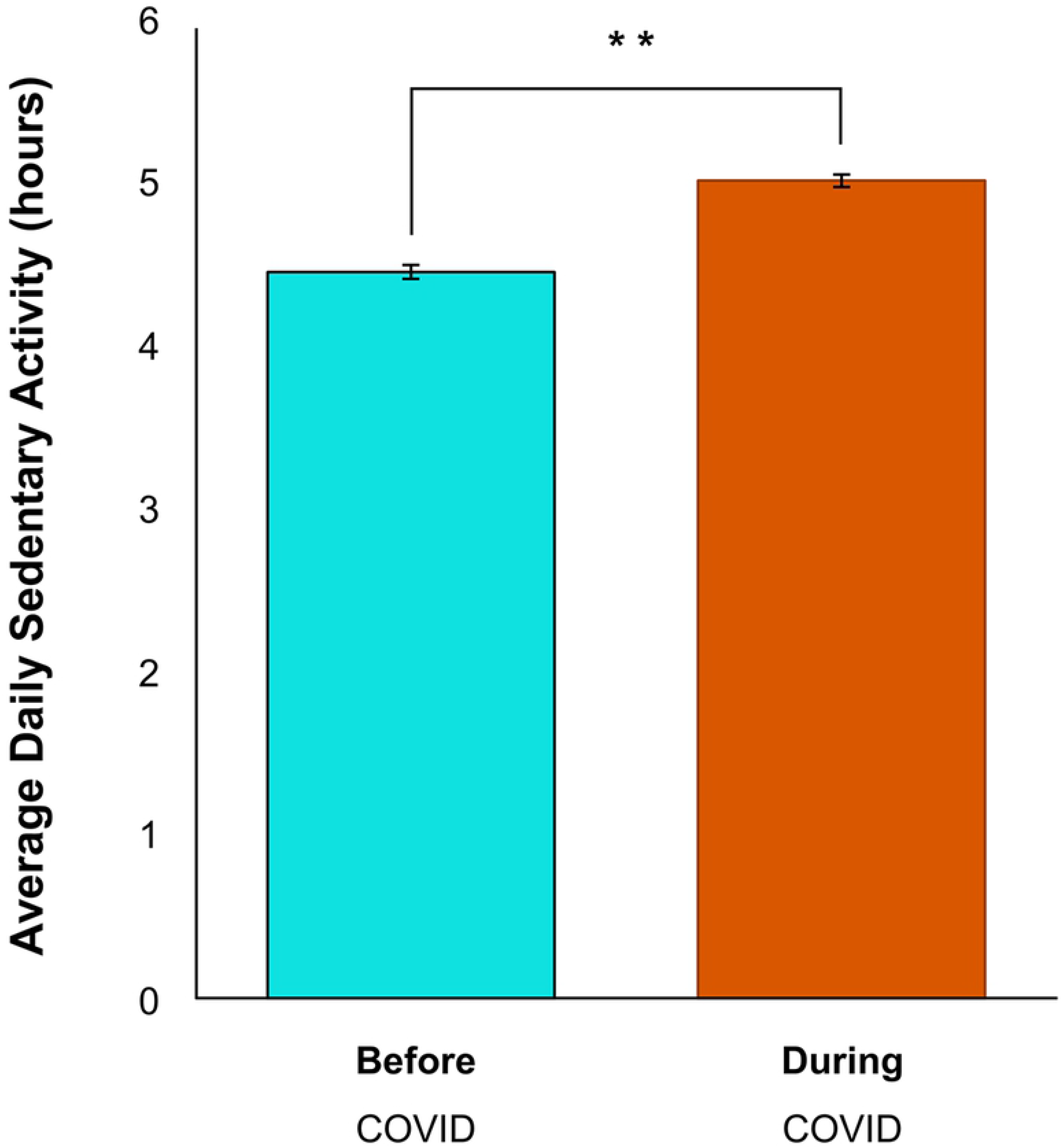
Average sedentary activity before and during the COVID-19 pandemic. There was a significant increase in sedentary activity reported by respondents since the COVID-19 pandemic (*Z* = −14.18, \*\**p* < 0.01). Error bars represent standard error.

**Fig 5.**
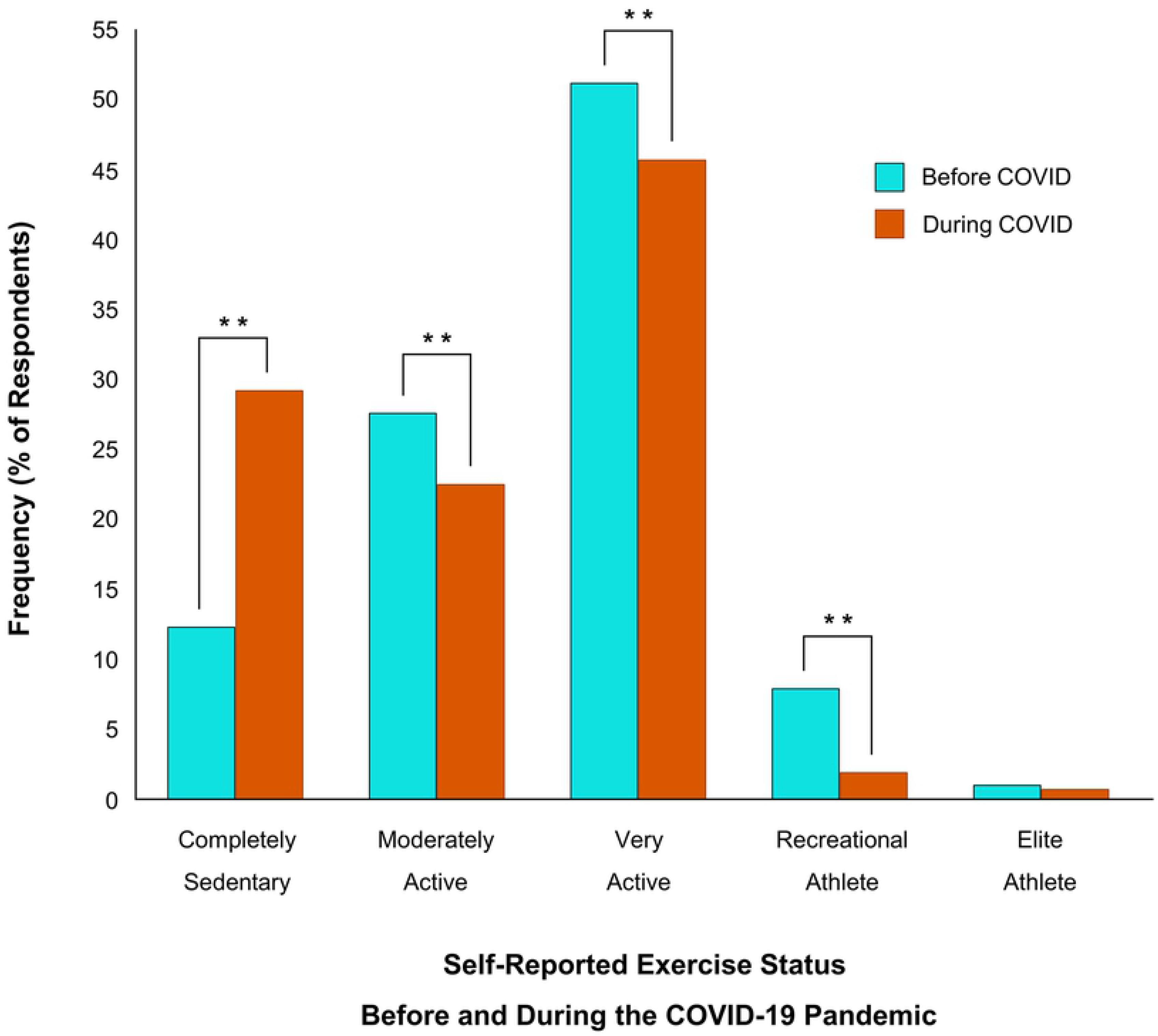
Self-report exercise status before and during the COVID-19 pandemic. 17% of respondents who had were “recreational athletes” (−6%; \*\**p* < 0.01), “very active” (−6%; \*\**p* < 0.01), or “moderately active” (−5%; \*\**p* < 0.01), now identify as being “completely sedentary” (\*\**p* < 0.01).

To identify respondents at higher risk for decreased physical activity during the initial stage of the COVID-19 pandemic, Kruskal-Wallis tests were conducted on between-group differences in physical activity change by income and age. Respondents who made “less than enough” (H(3,1384) = −3.60, *p* < 0.01) or “just enough” (H(3,1384) = −2.96, *p* < 0.01) income to meet their needs had significantly lower levels of MVPA during COVID-19 than those making “more than enough”. Although this trend was seen overall, the effect was largest for the 18-29 age group. A similar trend was observed in sedentary behaviour wherein those who made “less than enough” experienced greater increases in daily sedentary time compared to those who made “more than enough” (H(3,1422) = 95.14, *p* < 0.01), and those who made “just enough” also had elevated sedentary time compared to those who made “more than enough (H(3,1422) = 56.61, *p* < 0.05). Similarly, those aged 18-29 reported the greatest sedentary time during the COVID-19 pandemic compared to any other age group.

### Did physical activity and sedentary behaviour predict mental health during the pandemic?

When examining the change in total physical activity level split by the change in mental health status, respondents whose mental health got “worse” or “much worse” had greater reductions in physical activity since COVID-19 than those who experienced “no change” or got “better” or “much better” (Figure 6; H(5,1381) = 7.23, *p* < 0.01; H(5,1381) = 6.23, *p* < 0.01).

**Fig 6.**
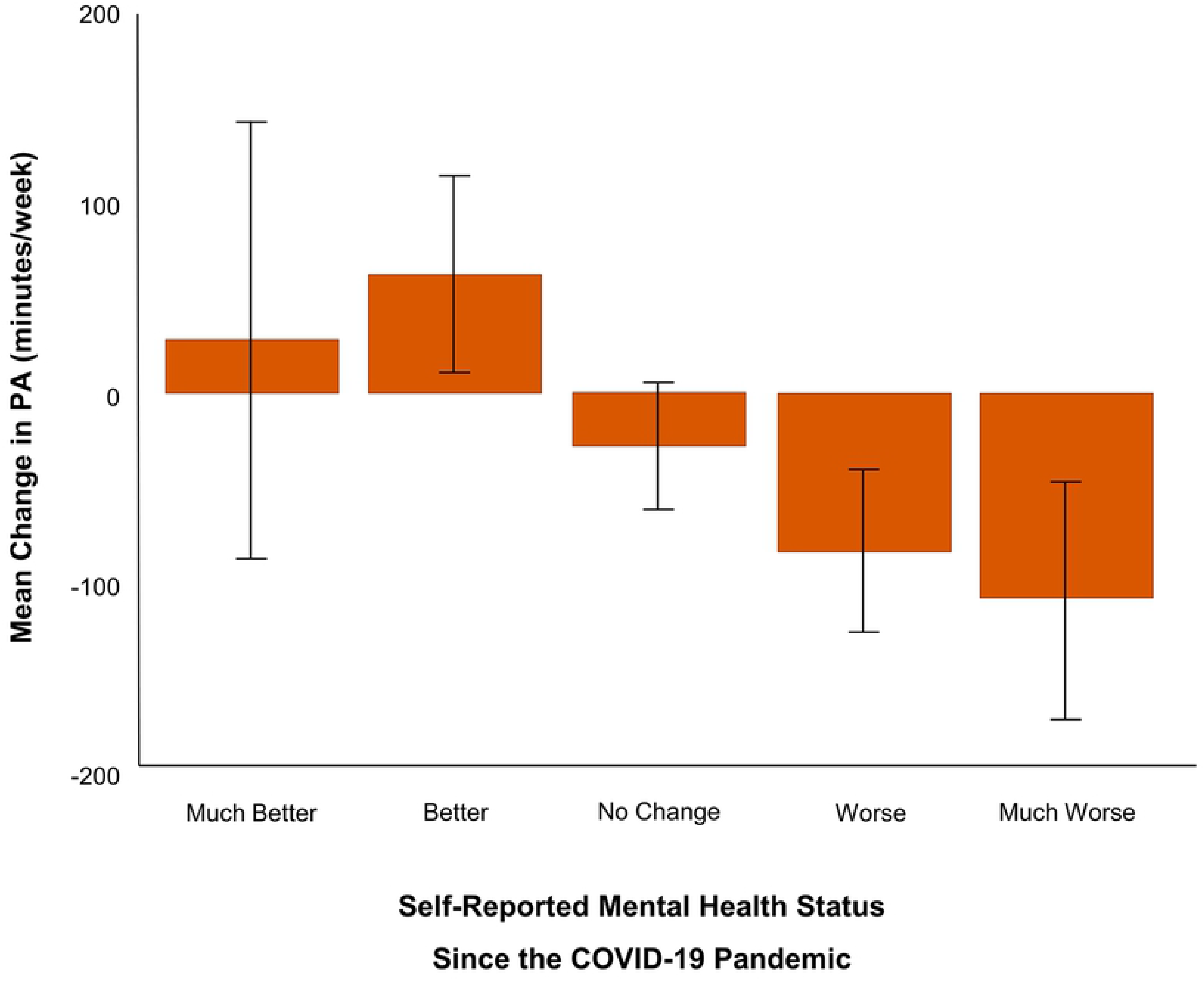
Change in total physical activity by change in mental health status. Respondents whose mental health got “worse” or “much worse” had greater reductions in physical activity time since COVID-19 compared to those who experienced “no change” or got “better” or “much better” (*p* < 0.01).

Spearman’s rank-order correlations were conducted assessing relationships between physical activity (prior, during, change) with mental health status (Tables 2 and 3). Overall, respondents who reported a greater decrease in their aerobic and strength-based physical activity during the pandemic also experienced more anxiety and depression (*r*(1544)= −0.12, *p* < 0.01; *r*(1544)= −0.21, *p* < 0.01). This was not only reflected their activity levels during the pandemic (i.e., people who engaged in less physical activity *during* the pandemic were more anxious and depressed; MVPA: *r*(1540)= −0.18, *p* < 0.01; *r*(1540)= −0.31, *p* < 0.01; ST: *r*(1542)= −0.14, *p* < 0.01; *r*(1542)= −0.22, *p* < 0.01); but also before (i.e., those who engaged in less physical activity *before* the pandemic were more depressed *during* the pandemic; *r*(1539)= −0.11, *p* < 0.01; *r*(1537)= −0.06, *p* < 0.01). Similar patterns in anxiety and depression were observed for those who experienced greater increases in sedentary behaviour, such that greater changes in sedentary time were associated with greater anxiety and depression during the pandemic (*r*(1420)= 0.07, *p* < 0.01; *r*(1420)= 0.12, *p* < 0.01).

### Impact of the pandemic on barriers and motivators to physical activity

To determine whether there were significant changes in the motivators and barriers to engage in physical activity due to COVID-19, McNemar’s tests were conducted. With respect to changes in motivators, respondents reported being less motivated to be physically active for ‘weight loss’ (−7%; *p* < 0.01), ‘strength building’ (−14%; *p* < 0.01), ‘enjoyment’ (−9%; *p* < 0.01), ‘appearance goals’ (down 4%; *p* < 0.01), ‘social engagement’ (−21%; *p* < 0.01), ‘sports training’ (down 5%; *p* < 0.01), and ‘healthcare provider recommended’ (−2%; *p* < 0.01). In contrast, respondents reported being more motivated to be physically active for ‘stress reduction’ (+5%; *p* < 0.01), ‘anxiety-relief’ (+14%; *p* < 0.01), ‘improve sleep’ (+4%; *p* < 0.05), and ‘no motivators’ (+4%; *p* < 0.01) (Figure 7). Viewing ‘increased energy’ as a motivator to engage in physical activity did not change during the pandemic (*p* > 0.05).

**Fig 7.**
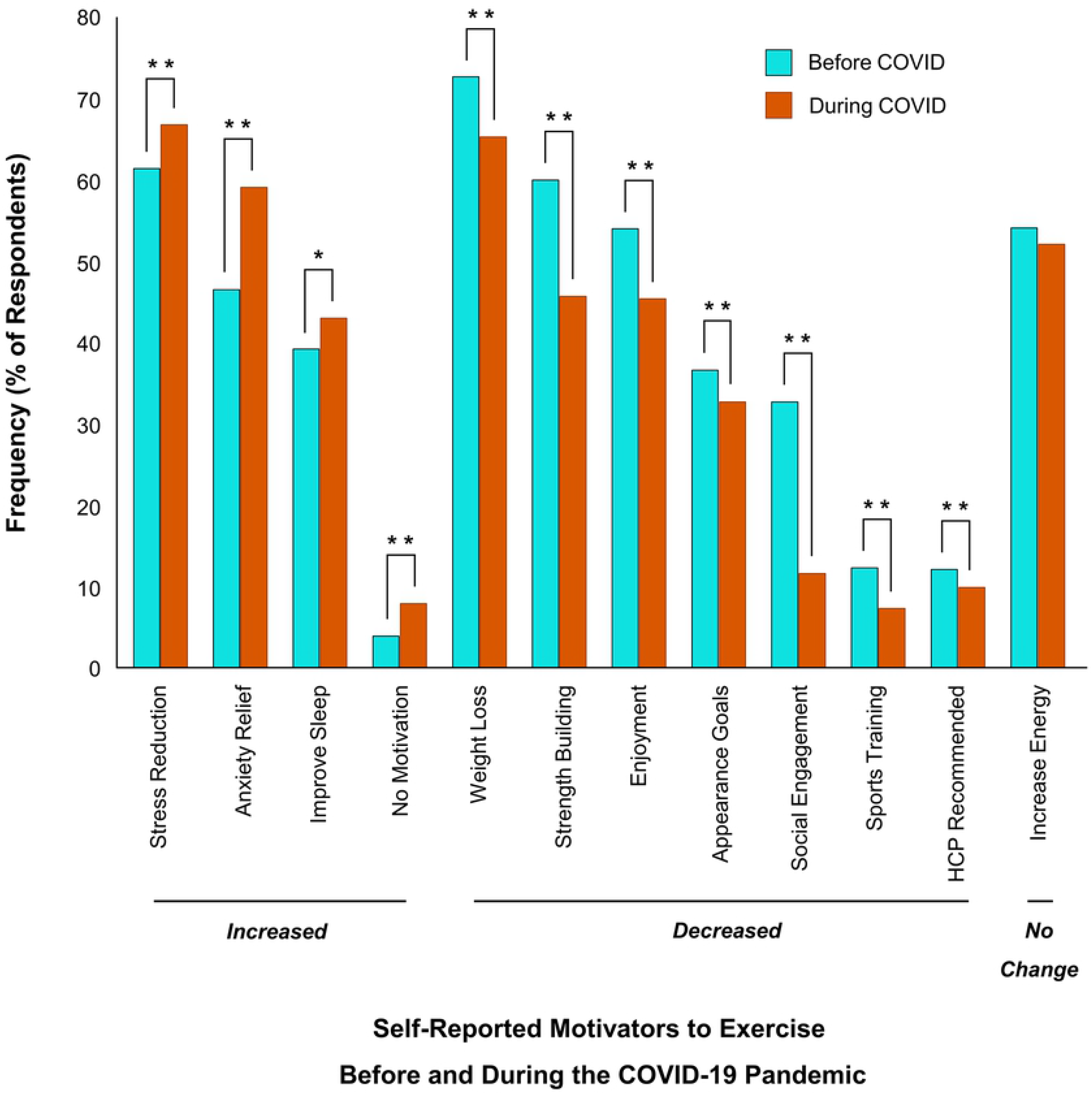
Changes in motivators to exercise before and during COVID-19. All motivators shown indicate a significant change (\**p* < 0.05, \*\**p* < 0.01). Motivators that increased significantly include ‘stress reduction’, ‘anxiety relief’, ‘improve sleep’ and ‘no motivators’. Motivators that decreased significantly include ‘weight loss’, ‘strength building’, ‘enjoyment’, ‘appearance goals’, ‘social engagement’, ‘sports training’ and ‘healthcare provider (HCP) recommended’. There was no change in how ‘increase energy’ was viewed as a motivator to exercise during the pandemic (*p* > 0.05).

With respect to barriers, respondents reported decreases in ‘insufficient time’ (−23%; *p* < 0.01), ‘no barriers’ (−10%; *p* < 0.01), ‘lack of confidence’ (−2%; *p* < 0.01), ‘recent injury’ (−3%; *p* < 0.01) and ‘insufficient finances’ (−3%; *p* < 0.01). In contrast, respondents reported increases in ‘lack of motivation’ (+8%; *p* < 0.01), ‘no facility access’ (+41%; *p* < 0.01), ‘no equipment’ (+23%; *p* < 0.01), ‘increased anxiety’ (+8%; *p* < 0.01), and ‘lack of support’ (+6%; *p* < 0.01). Barriers to engage in physical activity including ‘no access to childcare’, ‘lack of enjoyment’ and ‘fear of injury’ did not change because of the pandemic (Figure 8).

**Fig 8.**
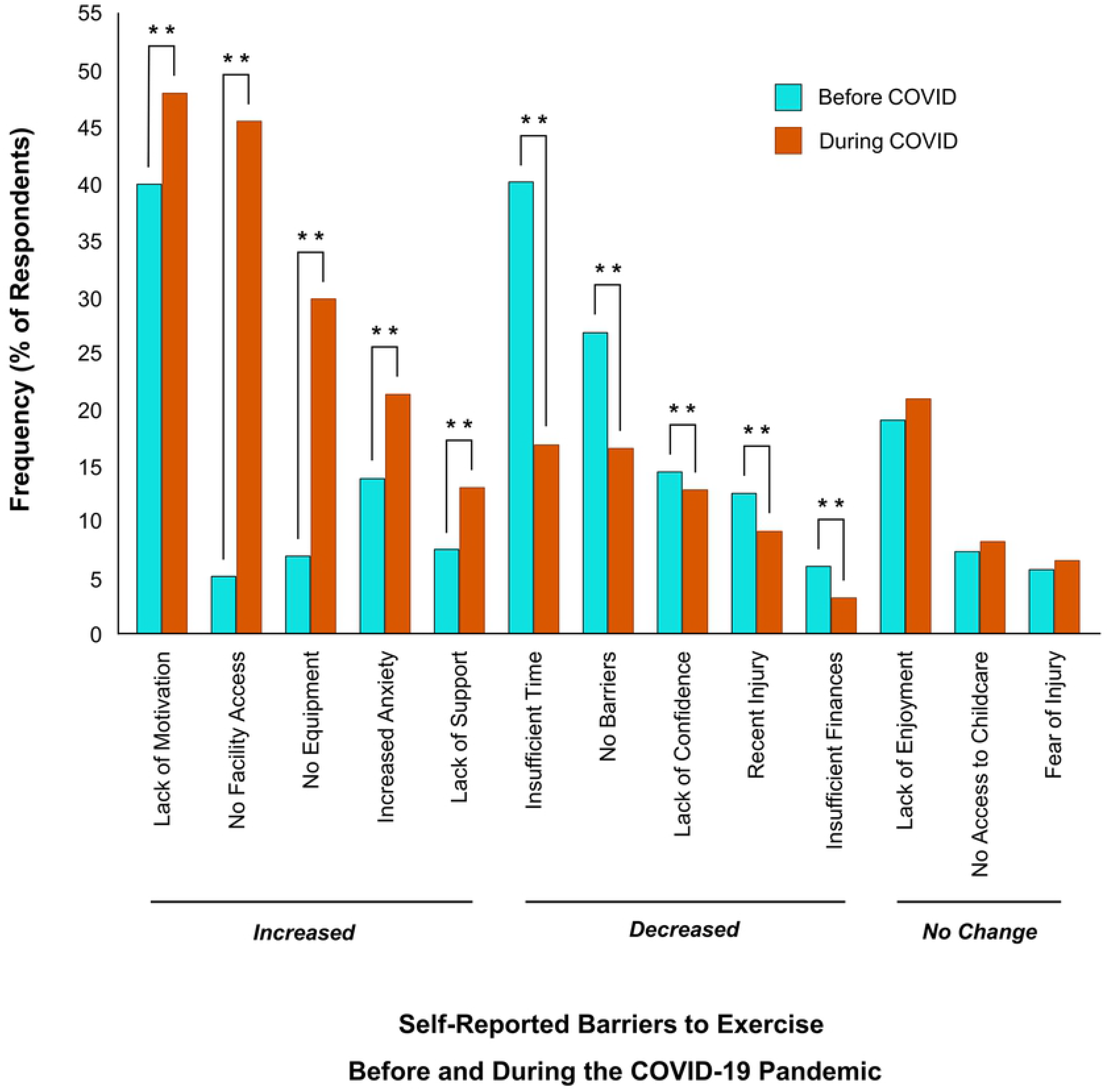
Changes in barriers to exercise before and during COVID-19. All barriers listed showed a significant change (\*\**p* < 0.01). Barriers which increased significantly since COVID-19 include ‘lack of motivation’, ‘no facility access’, ‘no equipment’, ‘increased anxiety’ and ‘lack of support’. Barriers which decreased significantly during COVID-19 include insufficient time’, ‘no barriers’, ‘lack of confidence’, ‘recent injury’ and ‘insufficient finances’. No change in barriers related to ‘lack of enjoyment’, ‘no access to childcare’ and ‘fear of injury’ (*p* > 0.05).

As an exploratory analysis, we conducted a series of linear regressions to determine whether self-reported levels of anxiety and depression predicted self-perceived barriers and motivators to exercise, and they did. Respondents who reported greater depressive symptoms were more likely to endorse ‘lack of self-motivation’ as a barrier to engaging in physical activity during the pandemic (*F*(1,1283) = 29.97, *p* < 0.01, R^2^= 0.02). Respondents who reported greater symptoms of anxiety were more likely to endorse ‘stress relief’ as a motivator (*F*(1,1282) = 26.05, *p* < 0.01, R^2^= 0.02) and ‘anxiety’ as a barrier (*F*(1,1283)= 7.16, *p* < 0.01, R^2^= 0.01) to engage in physical activity during the pandemic.

## Discussion

The present study examined the effect of the COVID-19 pandemic on the mental health, physical activity, and sedentary behavior of individuals undergoing pandemic lockdowns and physical distancing measures. Respondents reported higher psychological stress and moderate levels of anxiety and depression brought on by the pandemic. At the same time, the pandemic made it more difficult for them to be active, with aerobic activity down 11%, strength training down 30%, and sedentary time up 17%. Critically, respondents whose physical activity declined the most during the pandemic also experienced the worse mental health outcomes. Whereas, the respondents who maintained their physical activity levels, despite the pandemic, fared much better mentally.

Why was it so difficult for people to stay active during the pandemic? To address this important question, we assessed barriers and motivators to being physically active that may have changed during the pandemic. Overall, respondents were not motivated to be physically active because they felt too anxious and lacked social support. Respondents who were able to maintain their activity levels noticed a shift in what motivated them: they were less motivated by physical health and appearance, and more motivated by mental health and wellbeing. Stress relief, anxiety reduction, and sleep improvements were among the top motivators that increased during the pandemic, and indeed, research supports the use of physical activity for brain health [30] stress management [31] and sleep quality [32].

However, our results highlighted a paradox with mental health being both a motivator and barrier to physical activity. People wanted to be active to improve their mental health but found it difficult to be active due to their poor mental health. For example, despite the anxiolytic effects of exercise [30], respondents viewed their anxiety as a barrier to being physically active. Likewise, respondents who were more depressed were also less motivated to engage in physical activity, and amotivation is a symptom of depression itself. Although this is not a new challenge for clinicians whose depressed patients struggle to adhere to a prescribed exercise program [33], the stressfulness of the pandemic has made this a global issue that now must be considered when devising physical activity programs to support the mental wellbeing of citizens.

Was the drop in physical activity from the pandemic a cause or consequence of worsened mental health? Although this study cannot answer that question, it suggests the benefits of a two-pronged approach in promoting physical activity during stressful times that includes: 1) adopting a mode of physical activity that supports mental health, and 2) providing support to help minimize perceived psychological barriers to exercise [34]. For example, symptoms of anxiety may increase with high-intensity exercise and therefore moderate-intensity exercise might be preferable [35]. At the same time, to help overcome “feeling too anxious to exercise”, people should be encouraged to schedule their physical activity ahead of time in a calendar [36] to reduce feelings of uncertainty and decision fatigue that can aggravate their anxiety symptoms [37].

Not surprisingly, government-mandated closure to gyms and other recreational training facilities made it more difficult for people to be physically active. This was realized as a lack of necessary space and equipment during the pandemic reported as major barriers to being physically active. The pandemic forced a shift in doing everything at home but not everyone’s home is large enough or well-equipped to support their physical activity needs. Indeed, income level was predictive of activity level during the pandemic. People who reported “just enough” or “less than enough” income experienced greater decreases in physical activity and worsening mental health, especially younger adults aged 18 to 29 years old. Interestingly, these findings do not mirror the common trend that physical activity level declines with age [17] and instead, highlight a potential interaction between age and income that may reveal unique barriers to being physical activity. It is plausible that younger adults who typically work longer hours and earn less wage, are lacking both the time (e.g., due to long hours) and space (e.g., smaller dwelling) to meet physical activity goals. Outdoor activity could be a viable substitute [38], although this was not permitted in some countries during the pandemic [39]. Furthermore, increasing the number of repetitions performed during resistance training exercises can serve to adjust relative training intensity if lack of equipment is perceived as a barrier [40].

On top of being less active, our respondents reported spending significantly more time seated. The pandemic increased sedentary time by 10% or approximately 30 minutes per day. Although this may not seem like a lot, increasing sedentary time by just one hour has been associated with a 12% greater risk of mortality over a 6-year period [41]. But sedentary behavior is not only associated with poor physical health [42], it is also associated with poor mental health including lower perceived ratings of mental health and poorer quality of life [43]. Prolonged periods of sedentary behavior increase inflammatory markers [44] that may exacerbate symptoms of depression and anxiety [45]. Breaking up sedentary time with short frequent breaks (e.g., 1-2 minutes every half hour) may be sufficient to negate the negative health outcomes sedentary behaviour. Research shows that shorter frequent breaks are easier to adhere to than longer infrequent breaks [46] and can reduce sedentary behavior by more than 35 minutes per day, which would be enough to counteract the reported increase observed in this study.

Despite the valuable insights provided by this study; it is not without limitations. Our sample consisted mainly of young (18-29), highly educated (Bachelor’s degree or higher), female-identifying Canadian inhabitants which may limit the generalizability of the results. On average, our respondents were meeting the physical activity recommendations [16], which is not representative of the population at large. Moreover, a self-reported web-based survey was used to collect data and therefore response accuracy was unverifiable, and respondents required a device to access the internet; however, our large sample size would help minimize the impact of individual bias in reporting.

In conclusion, our findings highlight the importance of physical activity in mental health. During stressful times, like the COVID-19 pandemic, people are motivated to be physically active for their mental health but may be too anxious or depressed to partake. Our results point to the need for additional psychological supports to help people maintain their physical activity levels during stressful times in order to minimize the psychological burden of the pandemic and prevent the development of a mental health crisis.

## Acknowledgments

The authors wish to thank Samantha Choy for helping with data collection and management, and Paulina Rzeczkowska for helping create the tables and graphics.

---

**S1 Appendix. Barriers and motivators survey questions**.

